# A comprehensive map of human glucokinase variant activity

**DOI:** 10.1101/2022.05.04.490571

**Authors:** Sarah Gersing, Matteo Cagiada, Marinella Gebbia, Anette P. Gjesing, Atina G. Coté, Gireesh Seesankar, Roujia Li, Daniel Tabet, Amelie Stein, Anna L. Gloyn, Torben Hansen, Frederick P. Roth, Kresten Lindorff-Larsen, Rasmus Hartmann-Petersen

**Author notes:** Corresponding authors: F.P.R., K.L.-L., R.H.-P.

## Abstract

Glucokinase (GCK) regulates insulin secretion to maintain appropriate blood glucose levels. Sequence variants can alter GCK activity to cause hyperinsulinemic hypoglycemia (HH) or hyperglycemia associated with GCK-maturity-onset diabetes of the young (GCK-MODY), collectively affecting up to 10 million people worldwide. Patients with GCK-MODY are frequently misdiagnosed and treated unnecessarily. Genetic testing can prevent this but is hampered by the challenge of interpreting novel missense variants. Here we exploited a multiplexed yeast complementation assay to measure both hyper- and hypoactive GCK variation, capturing 97% of all possible missense and nonsense variants. Activity scores correlated with *in vitro* catalytic efficiency, fasting glucose levels in carriers of *GCK* variants and with evolutionary conservation. Hypoactive variants were concentrated at buried positions, near the active site, and at a region of known importance for GCK conformational dynamics. Some hyperactive variants shifted the conformational equilibrium towards the active state through a relative destabilization of the inactive conformation. Our comprehensive assessment of GCK variant activity promises to facilitate variant interpretation and diagnosis, expand our mechanistic understanding of hyperactive variants, and inform development of therapeutics targeting GCK.

## Introduction

Glucokinase (GCK) is the body’s primary glucose sensor as it regulates glucose-stimulated insulin secretion. Variants that decrease GCK activity cause elevated fasting glucose levels, known as GCK-maturity-onset diabetes of the young (GCK-MODY, MIM# 125851) (Froguel et al., 1992; Hattersley et al., 1992). Unlike most forms of diabetes, GCK-MODY does not require treatment, as glycemia remains unaltered (Stride et al., 2014), and patients do not suffer complications (Steele et al., 2014; Szopa et al., 2015). However, patients with GCK-MODY are often misdiagnosed with either type 1 or type 2 diabetes (Kleinberger et al., 2017; Pihoker et al., 2013; Shields et al., 2010), leading to unnecessary treatment and surveillance (Stride et al., 2014). Correct diagnosis of GCK-MODY can therefore terminate pharmaceutical treatments and decrease surveillance (Stride et al., 2014), with both economic and lifestyle benefits. Diagnosis of GCK-MODY can be achieved by identifying pathogenic variants in the gene encoding glucokinase (GCK, hexokinase-4).

GCK regulates glucose levels by catalyzing the first step of glycolysis — the phosphorylation of glucose to form glucose-6-phosphate. Glucose phosphorylation is the rate-limiting step in insulin secretion in pancreatic β-cells (German, 1993; Meglasson and Matschinsky, 1986; Meglassun et al., 1984) and glycogen synthesis in liver cells (Ferre et al., 1996), and these processes are therefore regulated by GCK activity.

In addition to GCK, there are three other human hexokinases. These hexokinases have a high affinity for glucose and show hyperbolic kinetics. In contrast, GCK has a low affinity for glucose (S_0.5_ 7.5-10 mM) (Vinuela et al., 1963) and sigmoidal kinetics, which together allow GCK to respond rapidly to changes in glucose levels in the physiological range. The sigmoidal kinetics of GCK are due to positive cooperativity with glucose (Hill coefficient 1.7) (Storer and Cornish Bowden, 1976). This positive cooperativity is unusual as GCK functions as a monomer and contains only one glucose binding-site. Instead, the sigmoidal response to glucose is caused by intrinsic protein conformational dynamics (Kamata et al., 2004; Larion et al., 2015; Lin and Neet, 1990).

GCK is a dynamic 52-kDa enzyme consisting of 465 amino acid residues, which fold into a large and a small domain. Between the two domains is a cleft forming the active site where glucose binds. The orientation of the two domains is not static, as GCK exists in multiple conformational ensembles (Larion et al., 2010; Sternisha et al., 2020). These ensembles are often described as three conformations: the super-open, open and closed conformation (Kamata et al., 2004). At low glucose levels, GCK predominantly exists in the super-open conformation. Upon glucose binding, GCK shifts into the open conformation (Liu et al., 2012; Zhang et al., 2006), and subsequently catalysis takes place in the closed conformation. Due to slow conversion between the super-open and open states (Heredia et al., 2006a), the population of each of the three conformations depends on glucose levels. At low glucose levels, GCK shifts into the super-open conformation prior to binding a new glucose molecule following catalysis, which results in slow glucose turnover. Turnover increases with higher glucose levels as GCK bypasses the super-open state and cycles between the open and closed conformations. The ratio between the slow and fast catalytic cycles, dependent on glucose concentration, gives rise to the positive cooperativity that is the basis for GCK sigmoidal kinetics and function (Kamata et al., 2004; Sternisha and Miller, 2019).

Underscoring the importance of GCK function for glucose homeostasis, variants that alter GCK activity are associated with several diseases (Osbak et al., 2009). Gain-of-function variants that increase GCK activity cluster at an allosteric activator site (Christesen et al., 2002; Gloyn et al., 2003) and cause hyperinsulinemic hypoglycemia (HH, MIM# 601820), which is characterized by increased insulin secretion even at low blood glucose levels (Christesen et al., 2002; Glaser et al., 1998). In contrast, loss-of-function variants where GCK activity is eliminated or decreased cause hyperglycemia. Inactivation of both GCK alleles can result in the severe permanent neonatal diabetes mellitus (PNDM, MIM# 606176) (Njølstad et al., 2001, 2003), while heterozygous mutations cause a mild form of diabetes known as GCK-MODY (Froguel et al., 1992; Hattersley et al., 1992). GCK-MODY has an estimated population prevalence of 0.11%-0.21% (Chakera et al., 2014; Ma et al., 2018), suggesting about 10 million people worldwide have GCK-MODY. Patients have mild and stable fasting hyperglycemia within 6-8 mM that does not usually require treatment, in contrast to other types of diabetes. Due to the population prevalence and as GCK-MODY is actionable, it has been proposed that GCK is suitable for inclusion in population screening programs (Schiabor Barrett et al., 2021). To act on variants detected in suspected GCK-MODY patients or in potential future population screening of GCK, we need to know the impact of any given variant on GCK function and its relation to diabetes risk. Without additional evidence, this will often not be possible: Of the 310 GCK missense variants reported in the ClinVar database (accessed on March 22nd 2022), 122 are currently classified as variants of uncertain significance (Landrum et al., 2020). Moreover, the scale of the challenge is immense. Given that every possible single-nucleotide change compatible with life already exists an average of 50 times in the human population (Weile and Roth, 2018), population screening will ultimately find all possible missense variants. To facilitate timely diagnosis, variant impacts would ideally be known proactively, in advance of each variant’s first presentation in a patient.

To address this challenge, we generated a variant effect map of GCK using multiplexed assays of variant effects (Fowler and Fields, 2014; Fowler et al., 2010). We assessed GCK activity using functional complementation in yeast, scoring both hypo- and hyperactive variants. The variant effect map recapitulates both the active site and a known allosteric activator site and includes 9003 of the 9280 (97%) possible missense and nonsense variants. The activity scores correlate with previous *in vitro* measurements of the catalytic efficiency and fasting blood glucose levels in patients. Furthermore, the map correctly classifies 77.9% of functionally characterized pathogenic variants that were previously curated (Osbak et al., 2009). To substantiate the map more broadly, we analyzed evolutionary conservation and conformational free energies. Conservation analysis generally agreed with the activity scores but was unable to capture hyperactive variants. When we examined these variants mechanistically, we found that some hyperactive variants likely shift GCK towards the active conformation through differential destabilization of the super-open and closed conformations. In conclusion, we present a comprehensive map of GCK activity to aid in variant interpretation and diagnosis of GCK-related diseases.

## Results

### Assessing human GCK activity using yeast complementation

To measure the activity of human GCK variants in large scale, we coupled yeast growth to human GCK activity using yeast complementation. To test complementation, we constructed a yeast strain deleted for all three yeast hexokinase genes (*hxk1Δ hxk2Δ glk1Δ*) that is unable to grow on glucose medium (Fig. 1A). This growth deficiency was rescued by expressing human pancreatic GCK (Fig. 1B), as previously shown (Mayordomo and Sanz, 2001).

**Fig. 1.**
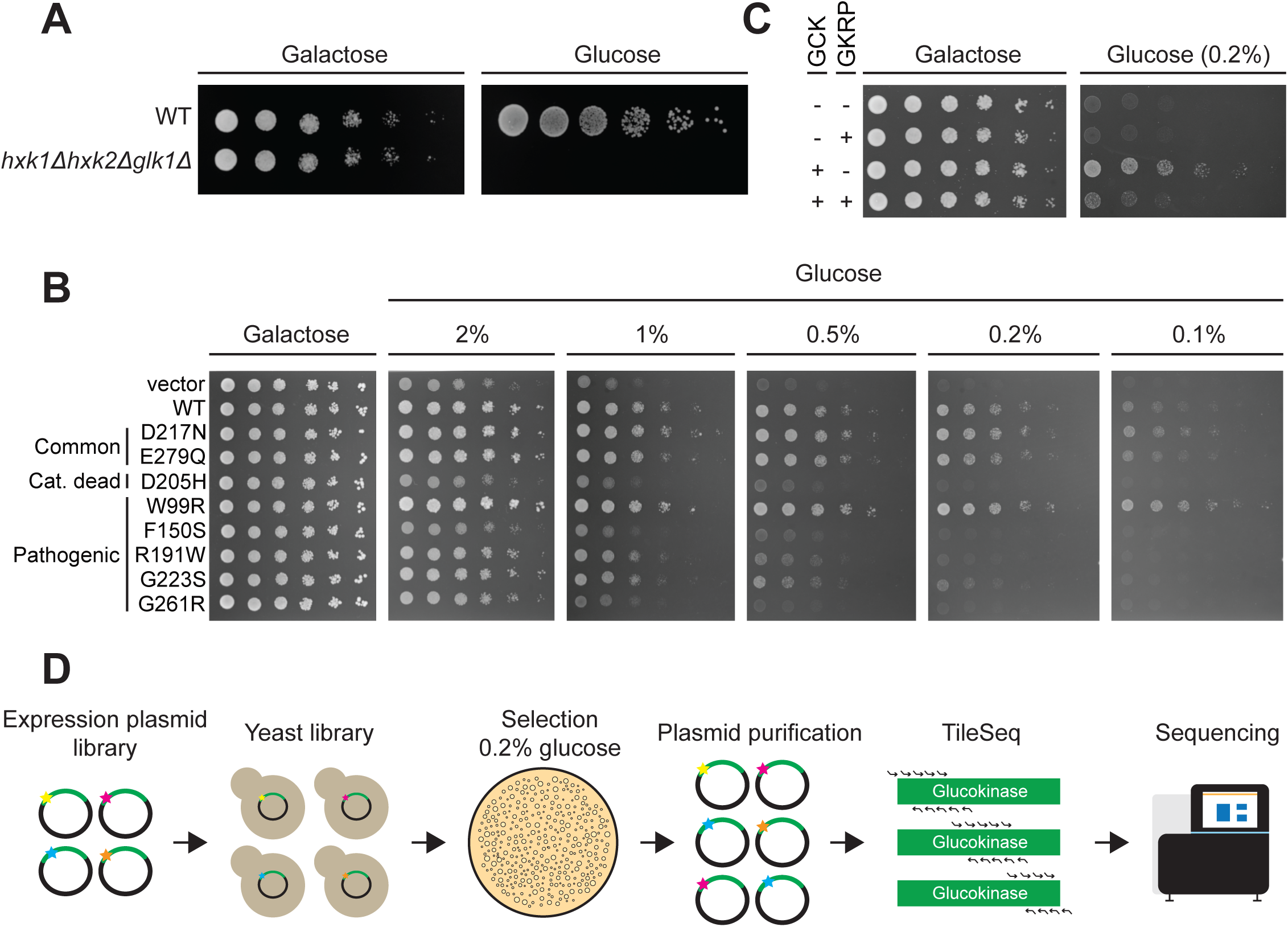
Yeast complementation as a readout for human glucokinase variant activity. (A) Yeast growth assay of wild-type (WT) and *hxk1Δ hxk2Δ glk1Δ* yeast strains on galactose and glucose media. (B) The growth of the *hxk1Δ hxk2Δ glk1Δ* yeast strain expressing either a vector control, wild-type GCK (WT) or a GCK variant was compared on media containing galactose or varying concentrations of glucose. (C) Growth assay of different combinations of vectors (-), GCK and GKRP expressed in the *hxk1Δ hxk2Δ glk1Δ* yeast strain on galactose and glucose media. (D) Illustration of the multiplexed assay for GCK variant activity.

Next, we examined whether the complementation assay could separate both hypo- and hyperactive pathogenic variants from non-pathogenic GCK variants. We selected an initial test set of eight variants. Two variants (p.D217N and p.E279Q) were used as non-damaging controls, as they were two of the three most common alleles in gnomAD (Karczewski et al., 2020), while a third variant (p.D205H) has previously been shown to be catalytically dead (García-Herrero et al., 2012; Kamata et al., 2004) and was used as a loss-of-function control. The five remaining test set variants were pathogenic variants from the ClinVar database (Landrum et al., 2020), including both one variant (p.W99R) associated with HH (Gloyn et al., 2003) and four variants (p.F150S, p.R191W, p.G223S and p.G261R) with compelling evidence for linkage to GCK-MODY (Ellard et al., 2000; Hattersley et al., 1998; Massa et al., 2001; Stoffel et al., 1992). To test complementation of the eight variants, they were expressed in the *hxk1Δ hxk2Δ glk1Δ* yeast strain (Fig. S1). On glucose medium, the two common variants showed growth that was similar to reference (‘WT’) GCK, while the catalytically dead variant (p.D205H) grew comparable to the vector control (Fig. 1B). The hyperactive variant (p.W99R) grew faster than WT, while the four disease-associated variants with reduced activity (p.F150S, p.R191W, p.G223S and p.G261R) grew slower than WT (Fig. 1B). The growth of the eight GCK variants was consistent with their level of activity, showing that the complementation assay could assess GCK activity.

GCK activity depends on the concentration of its substrate glucose. To find a glucose concentration that enabled detection of variants with both decreased and increased glucose affinities, we next tested complementation on media with varying glucose concentrations (Fig. 1B). A concentration of 0.2% (11.1 mM) glucose enabled detection of the hyperactive variant (p.W99R), while retaining a good dynamic range between WT and variants with decreased activity (p.F150S, p.R191W, p.G223S and p.G261R) (Fig. 1B). This concentration is close to GCK’s affinity for glucose (S_0.5_ = 7.5-10 mM) (Osbak et al., 2009; Vinuela et al., 1963).

To further establish the fidelity of the yeast system, we tested whether glucokinase regulatory protein (GKRP), a known inhibitor of hepatic GCK in humans (van Schaftingen, 1989; van Schaftingen et al., 1997), could repress GCK activity in the yeast system. We co-expressed human GKRP and GCK in the *hxk1Δ hxk2Δ glk1Δ* yeast strain and examined growth on glucose medium (Fig. 1C). Co-expression of GKRP on glucose medium lead to reduced growth (Fig. 1C). As GCK expression levels were comparable (Fig. S2), human GKRP inhibited GCK activity in yeast, further validating the relevance of the yeast system to assay human GCK activity.

Having established an assay coupling yeast growth to GCK enzymatic activity, we used the assay to score the activity of a saturated library of human GCK produced by codon-randomization. For library construction, we divided the GCK sequence into three regions that were separately mutagenized, assayed and sequenced. Regional libraries were mutagenized using oligos containing a central NNK-degenerate codon targeting each codon within individual regions. Subsequently, we cloned the mutagenized regional libraries *en masse* into a yeast expression vector and transformed the resulting plasmid libraries into the *hxk1Δ hxk2Δ glk1Δ* yeast strain. The yeast libraries were grown on 0.2% glucose media to select for GCK activity. Before and after selection, plasmids were extracted from yeast libraries and each region was deeply sequenced in tiles of ∼150 bp (∼1.6M-4.8M reads per sequenced tile) (Fig. 1D) (Weile et al., 2017). The resulting reads were used to quantify the relative frequency of each variant both before and after selection, and thus calculate an activity score and an associated measurement error for each variant. Activity scores were rescaled such that WT-like variants had a score of one while total loss-of-function variants had a score of zero. The resulting dataset contained scores for 9003 of the 9280 (97%) possible single amino acid GCK variants (including stop codons) (Fig. 2A), and most of the remaining variants could be imputed using the Human Protein Variant Effect Map Imputation Toolkit (Weile et al., 2017; Wu et al., 2019) (Fig. S3AB).

**Fig. 2.**
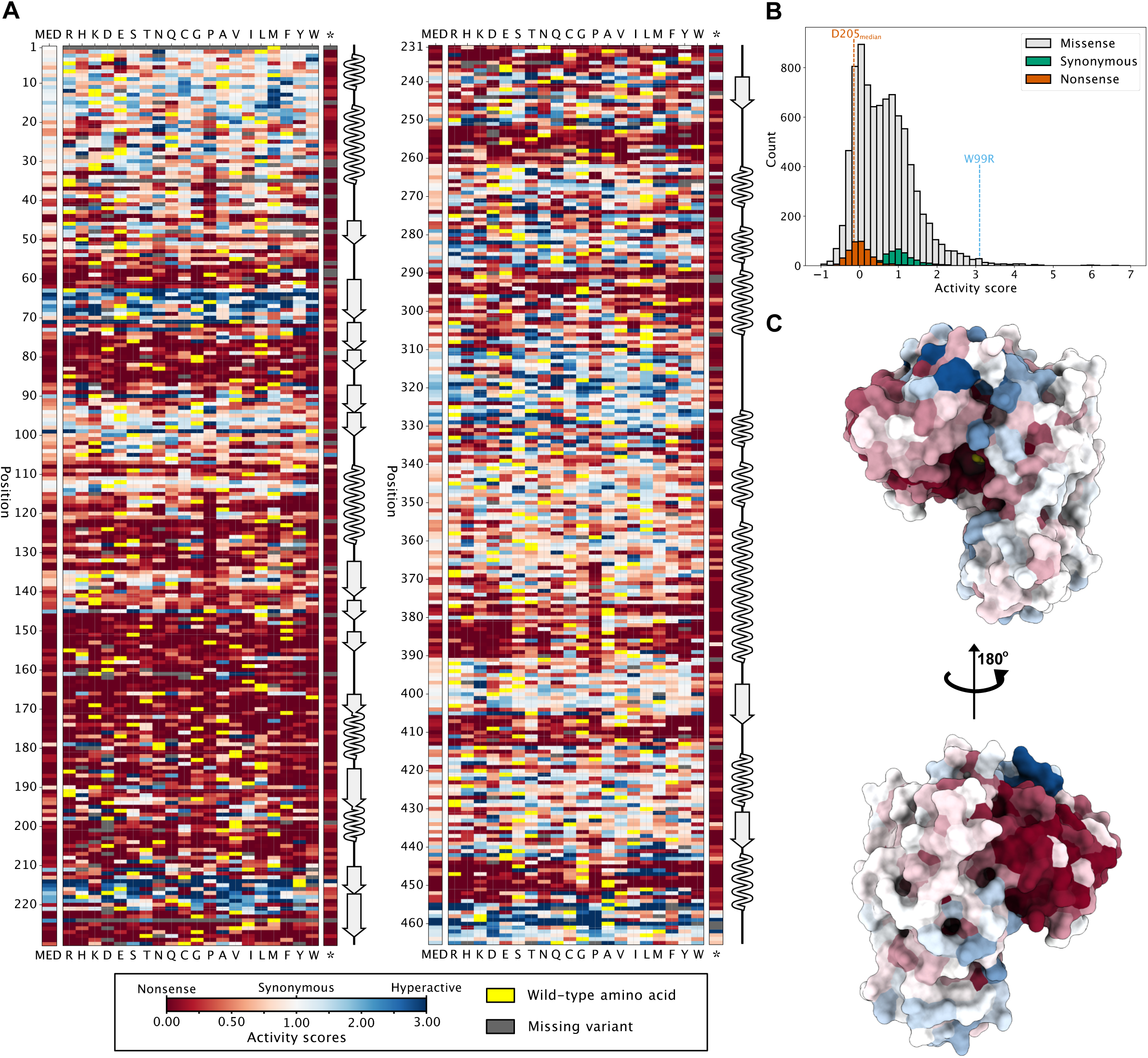
Map of glucokinase variant activity. (A) Heatmap showing the GCK activity score for each substitution (x-axis) at every position (y-axis) based on the multiplexed functional complementation assay. A score of one (white) corresponds to the activity of wild-type-like synonymous variants, a score of zero (red) corresponds to total loss-of-function and a score above one (blue) corresponds to activity above synonymous WT variants. The median (MED) activity score at each position is included. Yellow indicates the wild-type amino acid and missing variants are shown in grey. The secondary structure of GCK is shown next to the activity map with β-sheets shown as arrows and α-helices shown as waves. A map where missing variants have been imputed can be found in Fig. S3. (B) Distributions of activity scores for synonymous variants (green), nonsense variants (red) and missense variants (grey). The activity score of the hyperactive variant p.W99R (blue) and the median activity score of the catalytic residue p.D205 (red) are shown. (C) The median activity score at each position was mapped onto the closed glucose-bound conformation of GCK (PDB 1V4S). The color scheme is the same as in panel A.

The activity scores of nonsense and synonymous variants were well separated (Fig. 2B), and the distribution of missense variants spanned from total loss of function to increased activity (Fig. 2B). We used the standard deviation of synonymous variants (0.4) to set the thresholds for variants with decreased (score <0.6) and increased (score >1.4) activity. Using these thresholds, the assay identified 4442 (51.8%) variants with decreased activity and 1198 (14%) variants with increased activity, while the remaining 2930 (34.2%) variants had wild-type-like activities. The activity of our test-set variants was similar to the low-throughput growth assays, such that the common variants (p.D217N and p.E279Q) had wild-type-like activity, the HH-associated variant (p.W99R) had increased activity and the GCK-MODY-associated variants (p.F150S, p.R191W, p.G223S and p.G261R) had decreased activity. Although the catalytic site variant p.D205H was not included in the map, all other variants at this position showed decreased activity as expected.

To examine variant effects structurally, we mapped the median activity score at each position onto the structure of glucose-bound GCK (Kamata et al., 2004) (Fig. 2C, Fig. S4AB). The resulting activity-colored structure was consistent with characteristics of proteins in general and GCK specifically. First, surface residues were generally resistant to mutations, while active site and buried residues were mutation-sensitive (Fig. 2C, Fig. S4AB). Second, several positions where mutations on average increased activity clustered at a known allosteric activator site (Christesen et al., 2002). Together with the expected behavior of our test-set variants, these observations support that the map reflects human GCK activity.

### Correlations with enzyme kinetics, fasting blood glucose levels, and clinical genetics

To examine what aspects of GCK activity the map reflects, we examined the correlation between our assay scores and previously published kinetic parameters of 38 variants characterized *in vitro* (Gloyn et al., 2004). Assay scores correlated with the catalytic efficiency (k_cat_/S_0.5_) of GCK variants (Fig. 3A, *r_s_ =* 0.76, 95% CI [0.58, 0.88]), indicating that our yeast assay captures GCK catalytic efficiency with a dynamic range that includes both decreased and increased values. Having established that our assay recapitulates *in vitro* GCK activity, we next examined the correlation with measures of fasting glucose in carriers of GCK variants. Samples were obtained from four different Danish populations and included a population-based cohort (n=6,058 (men: 3,020; women: 3,038); Glümer et al., 2003), patients with newly diagnosed type-2 diabetes (n=2,855 (men: 1,678; women: 1,177); Christensen et al., 2018), a population of Danish children (n=1,138; (boys: 508; girls: 630); Kloppenborg et al., 2018) as well as ten patients diagnosed with GCK-MODY (men: 4; women: 6); Johansen et al., 2005). Our dataset included fasting glucose levels of 33 patients with known GCK variants and presenting with fasting glucose levels below 9 mM. Although fasting glucose levels likely depend on several genetic and environmental factors (Chen et al., 2021; Cuesta-Mũnoz et al., 2010; Dupuis et al., 2010; Simonis-Bik et al., 2008; Wędrychowicz et al., 2017), activity scores correlated with patient glucose levels (Fig. 3B, *r_s_* = -0.58, 95% CI [-0.20, -0.79]) suggesting that the yeast assay reflects the genetic contribution of GCK to fasting glucose in carriers of GCK variants. We next examined whether the scores of known GCK-MODY, HH, and benign variants separated into distinct classes. Our dataset included 71 variants with experimentally determined activity scores: 68 (60 GCK-MODY, 8 HH) pathogenic variants which have previously been functionally characterized (Osbak et al., 2009) and three benign variants (Beer et al., 2012; Landrum et al., 2020; Steele et al., 2011). We were unable to generate a score for the benign variant, p.G68D, and we therefore used the imputed score for this variant due to the already limited number of benign variants. Although the assay did not correctly classify all variants, variants belonging to each of the three classes showed distinct distributions centered on either scores comparable to synonymous mutations (benign variants), a hyperactive score (HH variants) or a decreased activity score (GCK-MODY variants) (Fig. 3C). Furthermore, variants with a high allele frequency (>10^-4^) in gnomAD (Karczewski et al., 2020) had WT-like scores while more rare variants displayed a wide range of activities (Fig. 3D). To determine threshold values to classify variants as either GCK-MODY or HH, we used receiver-operating characteristic (ROC) analyses, noting that the very small number of benign variants hampers a detailed analysis. The threshold for variants associated with GCK-MODY was 0.66 (AUC = 0.88), while variants with a score above 1.18 were predicted to be associated with HH (AUC = 0.94). Using these cutoff values, the activity assay correctly identifies 76.7% of the analyzed GCK-MODY variants and 87.5% of the analyzed HH variants.

**Fig. 3.**
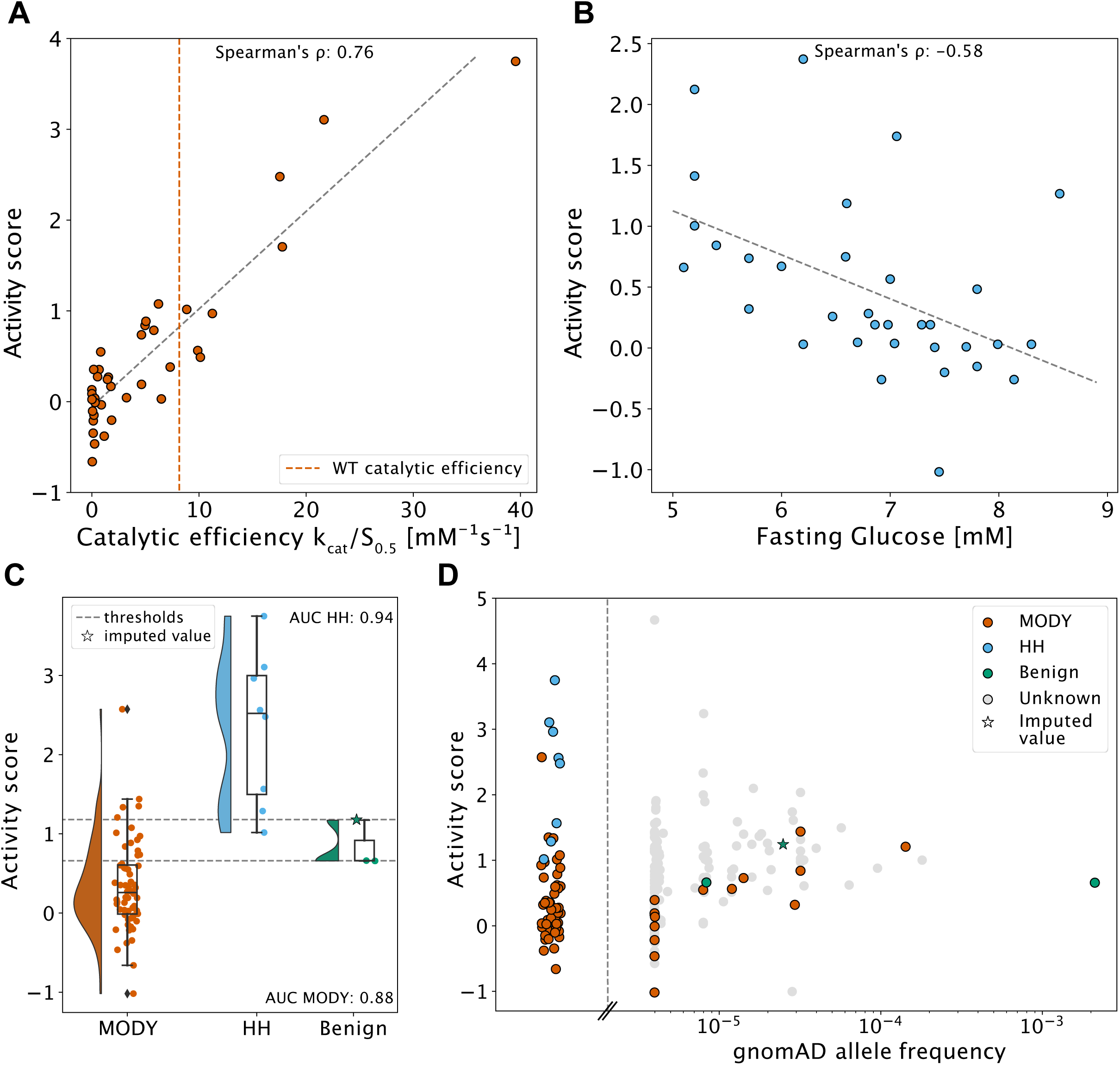
Correlations with enzyme kinetics, fasting blood glucose levels, and clinical genetics. (A) Plot showing the correlation between the GCK activity scores and previously measured catalytic efficiency (k_cat_/S_0.5_) (Gloyn et al., 2004) of 38 variants with a Spearmans’ ρ of 0.76. The red dotted line indicates WT catalytic efficiency (Gloyn et al., 2004) and the black dotted line shows the best fitting curve. (B) The activity score plotted against the fasting glucose level of 33 carriers with an identified single variant in the GCK gene. The black dotted line indicates the best fitting curve. Spearmans’ ρ is -0.58. (C) Raincloud plot showing the distributions of activity scores for 60 functionally characterized glucokinase-maturity-onset diabetes of the young (GCK-MODY) variants (Osbak et al., 2009), eight functionally characterized hyperinsulinemic hypoglycemia (HH) variants (Osbak et al., 2009) and three benign variants (Beer et al., 2012; Landrum et al., 2020; Steele et al., 2011). Due to the limited number of benign variants, the imputed activity score of the benign variant p.G68D was used, as no experimental score was obtained for this variant. The black dotted lines show the thresholds for variants associated with GCK-MODY (0.66, AUC = 0.88) and HH (1.18, AUC = 0.94) based on receiver-operating characteristic (ROC) analyses. (D) Plot showing the activity scores and the associated gnomAD allele frequency (Karczewski et al., 2020) for GCK variants present in gnomAD. As in panel C, the imputed activity score of p.G68D was used. In addition, the pathogenic variants from panel C that were not present in gnomAD are plotted to the left of the stippled line.

Although our assay was able to detect the vast majority of pathogenic variants, some reported pathogenic variants appeared as benign. There were several causes for this misclassification, but as examples we include three variants (p.V62M, p.T65I, and p.H137R) that show different molecular mechanisms causing them to score as benign in our assay. The genetic evidence for p.V62M as a loss-of-function mutation is compelling but initial *in vitro* kinetic characterization demonstrated increased affinity for glucose. Subsequent functional studies demonstrated that it is thermally labile with evidence for defective regulation by both GKRP and allosteric activators (Gloyn et al., 2005). Similarly, the HH variant p.T65I has an increased affinity for glucose (reduced S_0.5_) and therefore induces insulin secretion at inappropriate glucose levels (Gloyn et al., 2003). However, p.T65I also has a decreased turnover number, resulting in a WT-like catalytic efficiency (Gloyn et al., 2004), which is the aspect of GCK activity that our assay captures. Finally, the GCK-MODY variant p.H137R has been previously reported to have WT-like kinetics but mildly decreased thermal stability (Davis et al., 1999). Although our assay was not able to detect the p.H137R instability variant, it could detect severely unstable variants such as p.E300K (Burke et al., 1999). Thus, based on an evaluation of these three mutations, which display complex mechanisms for their pathogenicity, we note that our assay may not detect all pathogenic variants involving complex mechanisms, including modest instability, especially for those with WT-like intrinsic catalytic efficiency.

### Evolutionary conservation predicts glucokinase variant effects

While our experiments probe nearly all possible GCK variants, only a limited number of variants have previously been characterized experimentally. We therefore examined GCK activity scores more broadly by analyzing evolutionary conservation across species. Conservation analysis can assess the mutational tolerance of each position in a protein (Ng and Henikoff, 2001; Stone and Sidow, 2005). We analyzed GCK evolutionary conservation using a sequence alignment of homologous proteins, evaluating the evolutionary distance between the GCK WT sequence and the single mutant variant, using an alignment that included both hexokinases and glucokinases more widely. The resulting evolutionary distance score (ΔE) quantifies the likelihood of a given substitution (Fig. S5AB). Therefore, a score close to zero suggests that a substitution is compatible with function and does not affect the structural stability of the native conformations, while variants with a highly negative score are likely to be detrimental. Activity scores correlated weakly with ΔE (*r_p_ =* 0.44, 95% CI [0.43, 0.46], Fig. S6A). However, the correlation increased when we compared the two datasets using the residue median score (*r_p_ =* 0.64, 95% CI [0.57, 0.68], Fig. S6B), as averaging decreases noise in both datasets. Residue-averaged ΔE and activity scores agreed on regions where mutations severely decreased activity as well as regions that were tolerant towards mutations (Fig. 4A, Fig. S6E). However, the evolutionary conservation analysis did not detect gain-of-function positions nor the mutational sensitivity of the ∼150-200 region that is likely specific for GCK compared to other hexokinases (Fig. 4A, Fig. S6E). Residues in this region include the 151-179 loop that undergoes a disorder to order transition upon glucose binding (Larion et al., 2012). The glucose-induced conformational changes in GCK are the basis for the sigmoidal kinetics and low glucose affinity that distinguishes GCK from other hexokinases. Hence, the ∼150-200 region and gain-of-function variants are likely not captured by the evolutionary analysis due to the broadness of the multiple sequence alignment (MSA). When we restricted our analysis to include only residues with a median assay score below 1.18, correlation with ΔE further increased (*r_p_ =* 0.73, 95% CI [0.69, 0.77], Fig. S6CD). In conclusion, analysis of evolutionary conservation supports the activity assay. However, our conservation analysis does not include effects that are GCK-specific, such as substitutions that increase activity or affect conformational regulation.

**Fig. 4.**
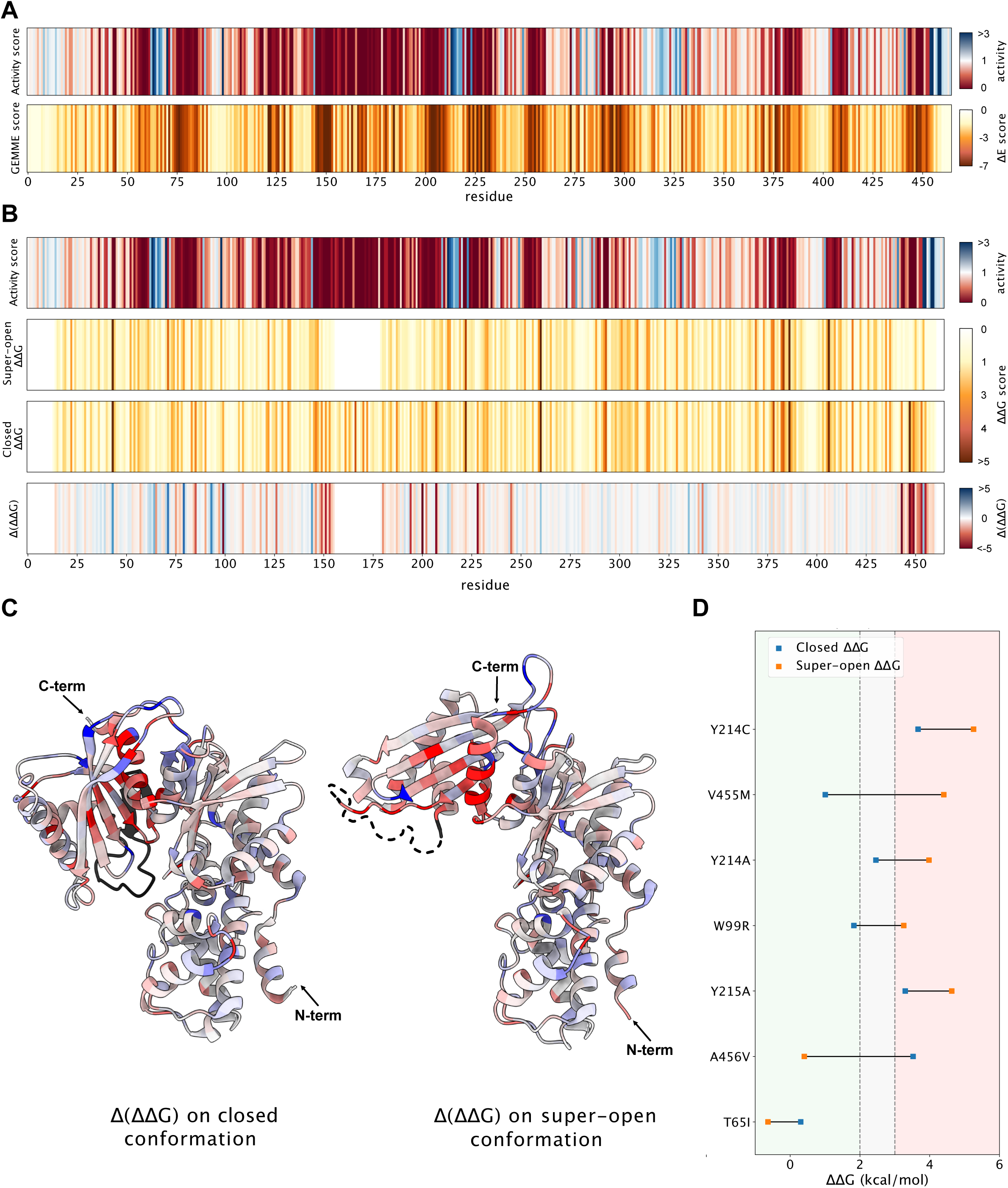
Interpreting the glucokinase activity scores using analyses of evolutionary conservation and conformational free energies. (A) Plots showing the residue median activity scores (top) and the residue median evolutionary conservation scores (GEMME score, bottom) for GCK. For activity scores, a score of one (white) corresponds to an activity comparable to wild-type-like synonymous variants, a score of zero (red) corresponds to total loss-of-function and a score above one indicates increased activity (blue). For GEMME scores (ΔE score), a score of zero (white) indicates that variants at this position are compatible with function, while a high negative score (red) indicates that variants at this position are detrimental. (B) Plots showing the residue median of activity scores, ΔΔG of the super-open conformation (PDB 1V4T), ΔΔG of the closed conformation (PDB 1V4S) and the difference between the ΔΔG in the closed and super-open conformation (ΔΔG_super-open_ – ΔΔG_closed,_ Δ(ΔΔG)), respectively. For activity scores, the color scheme is the same as in panel A. For ΔΔG of the super-open and closed conformations, a score of zero (white) corresponds to the same stability as wild-type, while a high negative score (red) means that variants at that position are less stable than wild-type GCK. For Δ(ΔΔG), a score of zero (white) indicates that variants at that position on average destabilize the super-open and closed conformations equally. A negative score (red) indicates that variants at that position on average destabilize the closed conformation relative to the super-open state, while a positive score (blue) indicates that variants at that position on average destabilize the super-open conformation relative to the closed state. Note that residues at positions 157-179 are absent from the structure of the super-open conformation, and scores at these positions are therefore not included in the plots of the super-open ΔΔG nor the Δ(ΔΔG). (C) The residue median Δ(ΔΔG) values from panel B are mapped onto the closed (PDB 1V4S) and super-open (PDB 1V4T) conformations of GCK. The 157-179 region absent from the super-open conformation is colored black in both structures. (D) Plot showing the ΔΔG values of seven previously characterized variants (Heredia et al., 2006b, 2006a) in the closed (PDB 1V4S) and super-open (PDB 1V4T) conformations. The background coloring indicates whether the ΔΔG value is in the stable (green), intermediate (grey) or unstable (red) range. Variants that are predicted to shift the equilibrium towards the closed conformation will have the highest ΔΔG in the super-open state (orange to the right). Conversely, variants that are predicted to shift the equilibrium towards the inactive state will have the highest ΔΔG in the closed conformation (blue to the right).

### Mechanistic evaluation of variant effects

To explore variants with functional impacts that were not detected via conservation analysis, we examined GCK variant effects mechanistically. We speculated that some variants shift the equilibrium of the conformational ensemble towards the catalytically inactive (or active) conformation, thereby decreasing (or increasing) GCK activity. This shift might arise if a given variant differentially affects the stability of the two conformations. To identify such variants we used Rosetta (Park et al., 2016) to predict changes in thermodynamic protein stability (ΔΔG) for both the super-open and closed GCK structure (Kamata et al., 2004) (Fig. S7AB), and then calculated the difference between the two structures (ΔΔG_super-open_ – ΔΔG_closed_, Fig. S7C). A negative score indicates that a given variant destabilizes the closed conformation relative to the super-open, while positive scores indicate variants that destabilize the super-open conformation relative to the closed. However, this shift is only relevant for variants that do not overly destabilize both structures, which would lead to a collapse of the enzyme structure, likely degradation and loss of function.

Variants at most positions destabilized the two conformations equally (Fig. 4B white lines, Fig. S8A). However, variants shifting the equilibrium towards the super-open inactive conformation were concentrated in two regions surrounding position 150 and 450 respectively (Fig. 4B lower panel, Fig. S8A). The region surrounding position 450 mapped to an α-helix that is surface-exposed in the super-open state but buried in the closed state (Fig. 4C), consistent with variants causing a greater destabilization of the closed state. Similarly, the residues surrounding position 150 were buried in the active conformation (Fig. 4C). In addition, this region is N-terminal to residues 151-179 that form a disordered loop in the super-open conformation and therefore are not included in the stability calculations. The disordered loop folds into a β-hairpin in the closed conformation (Kamata et al., 2004; Larion et al., 2012). Due to this disorder-order transition, mutations in the 151-179 loop/β-hairpin are likely to destabilize the closed state relative to the super-open state. A conformational shift towards the inactive state is therefore a potential mechanism for mutations in the ∼150-200 region. Positions where variants shifted the equilibrium towards the closed active conformation (Fig. 4B blue lines) were spread throughout the GCK sequence. However, these positions concentrated in the 3D structure at the allosteric activator site (Fig. 4C), which is also evident from the structure of GCK bound to a synthetic allosteric activator (Fig. S8BC). Previously, several hyperactive variants were examined to determine their mechanism using kinetic analysis (Heredia et al., 2006b, 2006a). By examining enzyme kinetics, Heredia *et al*. found that p.T65I, p.Y215A and p.A456V mainly accelerate glucose binding, while p.W99R, p.Y214A, p.Y214C and p.V455M predominantly increase the conformational preference towards the active state. Although p.T65I, p.Y215A and p.V455M affect the equilibrium mildly, all variants except p.A456V to some extent increased conversion to the closed state (Heredia et al., 2006b). Consistently, p.A456V shifted the equilibrium towards the super-open conformation according to our stability predictions (Fig. 4D). In addition, except for the p.T65I variant, stability calculations predicted the remaining five variants (p.W99R, p.Y214A, p.Y214C, p.Y215A and p.V455M) to shift the equilibrium towards the active conformation (Fig. 4D). Thus, for six out of seven variants our stability predictions were consistent with prior kinetic and mechanistic analysis, thereby validating our mechanistic predictions. We next used the stability calculations to assess how many hyperactive variants potentially shift the equilibrium towards the active state. Table 1 shows a list of 467 hyperactive variants (activity score > 1.18), which are predicted to both be stable (ΔΔG_closed_ < 2 kcal/mol) and promote increased populations of the closed active conformation (Δ(ΔΔG) > 0.25 kcal/mol).

## Discussion

Recent developments in molecular biology and sequencing technologies have made it possible to perform deep mutational scanning experiments, in which thousands of gene variants are assayed in a single, multiplexed experiment. Here we applied this technology to GCK, a gene of central importance in metabolism and where missense and nonsense variants are associated with several diseases. The resulting activity map covers 97% of all possible single amino acid variants, and activity scores correlate with *in vitro* catalytic efficiency, fasting glucose levels in individuals carrying GCK variants, and evolutionary conservation.

There are a number of limitations to our study. First, due to our mutagenesis strategy, each variant in the library may contain multiple mutations within the mutagenized region. As each regional library is sequenced in tiles, only *in cis* mutations that occur within the same ∼150 bp tile are detected. To limit the impact of potential background mutations that co-occur in the same clone(s), the regional libraries have a low mutational density, with an estimated average of 0.3-0.4 mutations per mutagenized region, and a high coverage (500,000x) so that each variant’s score is based on a large number of independent clones. In this way, the impact of secondary variants within a clone can be averaged out (just as genome-wide associations can associate traits with single variants despite genetic backgrounds that vary between individuals in the population). Indeed, TileSeq has previously been shown to perform on par with alternative strategies requiring full-length sequencing of each clone (Weile et al., 2017). A second limitation is that our mutagenesis libraries include only missense and nonsense variants. Other types of pathogenic variation have previously been identified in the GCK gene, including insertions (García-Herrero et al., 2007; Sayed et al., 2009), a promoter variant (Gašperíková et al., 2009), and splice site variants. A map including only missense and nonsense variants is, however, still highly relevant. First, the effects of missense variants can be challenging to interpret without functional data. Second, missense and nonsense variants constitute 335 out of the total 566 current GCK ClinVar entries (Landrum et al., 2020) (accessed on March 22^nd^ 2022). Finally, of the 184 GCK variants that have been annotated as variants of uncertain significance, 122 (66%) of these are missense variants. A third limitation of our study is that 22.1% of the analyzed pathogenic variants have scores that are similar to those of wild-type-like synonymous variants. The sensitivity of the assay could potentially be increased by repeating the assay using different expression levels and glucose concentrations as well as increasing the number of replicates. However, others have found that pathogenic variants, even when thoroughly examined in low-throughput assays, can appear wild-type-like (Gloyn et al., 2004; Miller et al., 1999; Sagen et al., 2006). Multiple assays characterizing different aspects of GCK function might therefore be necessary to capture all pathogenic variants.

The hyperactive variants identified in our assay clustered at a site distal from the active site, known as the allosteric activator site (Christesen et al., 2002). The allosteric site is being pursued as a target for treatment of type 2 diabetes using drugs known as glucokinase activators (GKAs) (Grimsby et al., 2003; Matschinsky, 2009). The development of GKAs has faced several problematic effects including accumulation of triglycerides in the liver (De Ceuninck et al., 2013; Rees and Gloyn, 2013) and hypoglycemia (Bonadonna et al., 2010; Meininger et al., 2011). However, recently both a dual-acting and a hepatoselective GKA have shown promising results (Vella et al., 2019; Zhu et al., 2018). Most GKAs bind the allosteric activator site and mimic the effect of known hyperactive variants. Previously, a few hyperactive variants were found to increase GCK activity by favoring the conformational change to the closed active state (Heredia et al., 2006b, 2006a). We extended the mechanistic analysis of hyperactive variants using predictions of protein stability for the super-open and closed conformations. Our findings are in accordance with prior kinetic analyses (Heredia et al., 2006b, 2006a) and we identify 467 hyperactive variants that are predicted to shift GCK towards the closed state. Furthermore, our results indicate that these variants facilitate isomerization by destabilizing the super-open conformation relative to the closed. In conclusion, our study provides a comprehensive mapping of the allosteric activator site and an improved mechanistic understanding of some hyperactive variants. Our results may aid in refining drug design and the development of GKAs, specifically to address the problem of hypoglycemia resulting from GKA treatment.

In contrast to hyperactive variants we found that, in the region spanning residues ∼150-210, nearly all substitutions substantially decreased activity. Variants previously identified in the region are associated with loss of activity and elevated fasting plasma glucose levels (Gloyn et al., 2004; Osbak et al., 2009), and this region is central in the conformational dynamics of GCK (Larion et al., 2012, 2015). Hence, substitutions in this region might interfere with the transitions required for activity. In addition, some variants might destabilize the closed active conformation relative to the super-open state, thereby leading to increased population of the inactive state. However, the mechanisms underlying the mutational sensitivity of this region are not entirely clear and require further studies.

Here, we provide the first comprehensive map of GCK variant activity, measuring the functional consequences of thousands of previously uncharacterized GCK variants. More than 1 in 1000 people are estimated to suffer from GCK-MODY, and although the functional evidence from our variant effect map cannot alone classify a variant as pathogenic, it can aid variant interpretation to diagnose GCK-MODY and reduce unnecessary treatment resulting from misdiagnosis.

## Materials and methods

### Buffers

TE Buffer: 10 mM Tris-HCl, 1 mM EDTA, pH 8.0. SDS sample buffer (4x): 250 mM Tris/HCl, 8% SDS, 40% glycerol, 0.05% bromophenol blue, 0.05% pyronin G, 2% β-mercaptoethanol, pH 6.8.

### Plasmids

The pancreatic isoform of human GCK (Ensembl ENST00000403799.8) was codon optimized for yeast expression and cloned into pDONR221 (Genscript). The initial test set GCK variants were generated by Genscript. For yeast expression, WT GCK, test set variants and libraries were cloned into pAG416GPD-EGFP-ccdB (Addgene plasmid # 14316 ; http://n2t.net/addgene:14316 ; RRID:Addgene_14316, (Alberti et al., 2007)) using Gateway cloning (Invitrogen). Human GKRP (Ensembl ENST00000264717.7) was codon optimized for yeast expression and cloned into pDONR221 with an N-terminal HA-tag (Genscript). For yeast expression, GKRP was cloned into pAG415GPD-ccdB (Addgene plasmid # 14146 ; http://n2t.net/addgene:14146 ; RRID:Addgene_14146, (Alberti et al., 2007)) using Gateway cloning (Invitrogen).

### Yeast strains

The *hxk1Δ hxk2Δ glk1Δ* strain used for GCK complementation assays was obtained in two steps. First, the following two strains were crossed to obtain a haploid *hxk1Δ hxk2Δ MATa* strain: *hxk1*::kanMX *his3Δ1 leu2Δ0 met15Δ0 ura3Δ0 MATa* and *hxk2*::natMX4 *can1Δ*::STE2pr-Sp_HIS5 *lyp1Δ0 his3Δ1 leu2Δ0 ura3Δ0* met15Δ0 LYS2+ MATα. This strain was then used to knock out GLK1 using a HygroMX cassette. BY4741 was used as a wild-type control. Yeast cells were cultured in synthetic complete (SC) medium (2% D-galactose, 0.67% yeast nitrogen base without amino acids, 0.2% drop out (Sigma), (0.0076% uracil, 2% agar)). To select for GCK activity, yeast cells were grown on SC medium containing various concentrations of D-glucose monohydrate as indicated in figures. The growth defect of the *hxk1Δ hxk2Δ glk1Δ* strain was tested on Yeast Extract-Peptone (YP) medium (2% D-glucose or 2% D-galactose, 2% tryptone, 1% yeast extract). Small scale yeast transformations were done as described in (Gietz and Schiestl, 2007a).

### Yeast growth assays

For yeast growth assays on solid medium, cultures were grown overnight (30 °C, vigorous agitation) until reaching exponential phase. The yeast cells were washed once in sterile water (1200 g, 5 min, RT) and resuspended in sterile water. The OD_600nm_ of all cultures were adjusted to 0.4, and the cultures were used for a five-fold serial dilution in water. Serial-diluted cultures were spotted in 5 µL drops onto agar plates, which were briefly dried and incubated at 30 °C for two to four days.

### Protein extraction from yeast cells

For extraction of proteins from yeast, cultures were grown overnight (30 °C, vigorous agitation) until reaching exponential phase, at which point 100-125 × 10^6^ cells were harvested (1200 g, 5 min, 4 °C). The pelleted cells were washed in 25 mL ice-cold water (1200 g, 5 min, 4 °C). The cell-pellet was resuspended in 1 mL 20% ice-cold TCA, transferred to an Eppendorf tube and centrifuged (4000 g, 5 min, 4 °C). The supernatant was discarded and the pellet was resuspended in 200 µL 20% ice-cold TCA. The resuspension was transferred to a screw cap tube containing 0.5 mL glass beads (Sigma), and cells were lysed using a Mini Bead Beater (BioSpec Products) by three 15 second cycles with 5 minute incubations on ice between each burst. Then, 400 µL ice-cold 5% TCA was added, the tubes were punctured at the bottom using a needle and transferred to a 15 mL Falcon tube containing a 1.5 mL Eppendorf tube without the lid. The sample was isolated from the glass beads by centrifugation (1000 g, 5 min, 4 °C). The Eppendorf tube containing the sample was centrifuged (10000 g, 5 min, 4 °C) and the resulting pellet was washed in 500 µL 80% acetone (10000 g, 5 min, 4 °C). The acetone was removed and the pellet was dried for 5 min before resuspension in 100 µL SDS sample buffer (1.5x) and 25 µL 1 M Tris/HCl, pH 9. The samples were boiled for 5 min and cleared by centrifugation (5000 g, 5 min, RT). The supernatant was transferred to an Eppendorf tube and was analysed by SDS-PAGE and Western blotting.

### Electrophoresis and blotting

SDS-PAGE was done using 12.5% acrylamide gels. Following SDS-PAGE, 0.2 µm nitrocellulose membranes were used for the Western blotting procedure. After protein transfer, membranes were blocked in 5% fat-free milk powder, 5 mM NaN_3_ and 0.1% Tween-20 in PBS. Antibodies and their sources were: anti-HA (Roche, 15645900) and anti-GFP (Chromotek, 3H9 3h9-100). The secondary antibody was HRP-anti-rat (Invitrogen, 31470).

### Library mutagenesis

To construct a library of GCK variants, oligos containing a central degenerate NNK codon were designed for each codon in the GCK sequence. An online tool (http://llama.mshri.on.ca/cgi/popcodeSuite/main) (Weile et al., 2017) was used to obtain oligo sequences, and oligos were obtained from Eurofins. The GCK sequence was divided into three regions spanning aa 2-171 (region 1), 172-337 (region 2) and 338-466 (region 3). Oligos were pooled for each region. The three regional oligo pools were phosphorylated using T4 Polynucleotide Kinase (NEB) as recommended by the provider using 300 pmoles of each oligo pool and incubation for one hour at 37 °C. Then, the phosphorylated oligos were annealed to the WT GCK sequence. For each region, the following was combined in a PCR tube: 25 fmoles pENTR221-GCK, 3 µL 10 µM SKG_1, 5.6 µL oligo pool, 10.1 µL nuclease-free water. The reactions were denatured (95 °C, 3 min) and cooled (4 °C, 5 min). Following template annealing, 5 µL of each reaction was combined with 5 µL Phusion Hot Start Flex 2x Master Mix (NEB) and sequences were extended (95 °C 3 min, 4 °C 5 min, 50 °C 120 min). To each reaction, the following was added: 1.5 µL Taq DNA ligase buffer (NEB), 0.5 µL Taq Ligase (NEB), 3 µL nuclease-free water, followed by incubation (45 °C 20 min). Next, 1 µL of each reaction was combined with the following: 2 µL 10 µM SKG_2, 2 µL 10 µM SKG_3, 25 µL Phusion High-Fidelity PCR Master Mix with HF Buffer (NEB), 20 µL nuclease-free water. The libraries were amplified using the following conditions: 98 °C 30 sec, 20 cycles of 98 °C 15 sec, 55 °C 30 sec, 72 °C 150 sec, followed by 72 °C 5 min and 4 °C hold. The resulting PCR products were used in a PCR to add Gateway attB sites: 25 µL Phusion High-Fidelity PCR Master Mix with HF Buffer (NEB), 1 µL PCR product, 2 µL 10 µM SKG_4, 2 µL 10 µM SKG_5, 20 µL nuclease-free water. The following PCR program was used: 98 °C 30 sec, 5 cycles of 98 °C 15 sec, 58 °C 30 sec, 72 °C 150 sec, followed by 12 cycles of 98 °C 15 sec, 72 °C 150 sec and finally 72 °C 5 min, 4 °C hold. The resulting PCR products containing Gateway attB-sites were gel purified and used for Gateway cloning.

### Library cloning

Next, PCR products were cloned into pDONR221 to generate three regional pENTR221 libraries. For each of the three regions a 25 µL Gateway BP reaction was prepared: 114 ng PCR product, 375 ng pDONR221, 5 µL Gateway BP Clonase II enzyme mix (ThermoFisher), TE Buffer pH 8.0 to 25 µL. Reactions were incubated overnight at RT. The following day, 3.1 µL proteinase K was added and reactions were incubated for 10 min at 37 °C. For each region, 4 µL BP reaction was transformed into 100 µL of NEB 10-beta electrocompetent *E. coli* cells using electroporation. The cells were recovered in 3900 µL NEB 10-beta outgrowth medium in 50 mL Falcon tubes at 37 °C for 1 hour. Then, cells were plated on LB containing kanamycin and were incubated at 37 °C overnight. A minimum of 500,000 colonies were obtained for each region. The following day, the cells were scraped from the plates using sterile water and plasmid DNA was extracted from 400 OD_600nm_ units (Nucleobond Xtra Midiprep Kit, Macherey-Nagel).

The resulting pENTR221-GCK libraries were used in large-scale Gateway LR reactions to clone the libraries into the pAG416GPD-EGFP vector. For each region, the following was mixed: 216.9 ng pENTR221-GCK library, 450 ng pAG416GPD-EGFP vector, 6 µL Gateway LR Clonase II enzyme mix (ThermoFisher), TE Buffer pH 8.0 to 30 µL. The reactions were incubated at RT overnight. Next day, 3 µL proteinase K was added to each reaction and tubes were incubated at 37 °C for 10 min. The LR reactions were transformed into NEB 10-beta electrocompetent *E. coli* cells using electroporation of 4 µL reaction per 100 µL cells. Cells were recovered in NEB 10-beta outgrowth medium for 1 hour at 37 °C, and were then plated on LB containing ampicillin for incubation at 37 °C overnight. A minimum of 500,000 colonies were obtained for each regional library. The following day, cells were scraped from plates using sterile water and plasmid DNA was extracted from 400 OD_600nm_ units (Nucleobond Xtra Midiprep Kit, Macherey-Nagel). The resulting plasmid DNA was used for yeast transformation.

### Library yeast transformation

The GCK expression libraries were transformed into the *hxk1Δ hxk2Δ glk1Δ* strain as described in (Gietz and Schiestl, 2007b) using a 30x scale-up and 30-60 µg of plasmid DNA. Small aliquots of each transformation were plated on SC-URA galactose in duplicate for colony counting. The rest of each transformation was diluted in SC-URA galactose to an OD_600nm_ of 0.2, and incubated at 30 °C with shaking until saturated. A minimum of 500,000 colonies were obtained for each regional library. For each regional library, 36 OD_600nm_ units of yeast cells were harvested in duplicate to serve as pre-selection samples. Pellets were stored at -20 °C before plasmid DNA extraction. In parallel with library transformations, a vector (pAG416GPD-EGFP-ccdB) and GCK WT control were transformed in small scale, and 36 OD_600nm_ units in duplicate were stored at -20 °C for DNA extraction.

### Library selection

Yeast transformations were next used for selection of GCK activity. For each regional library, 20 OD_600nm_ units of yeast cells were harvested in duplicate. The cells were washed three times in sterile water and resuspended in 500 µL sterile water. The cells were then plated on large plates (500 cm^2^) of SC-URA 0.2% glucose. Plates were incubated at 30 °C for three days. After selection, yeast cells were scraped off plates using 30 mL milliQ water and 36 OD_600nm_ units from each plate were harvested to serve as post-selection samples. Cell pellets were stored at -20 °C prior to plasmid DNA extraction. In parallel 2.6 OD_600nm_ units of vector (pAG416GPD-EGFP-ccdB) and GCK WT yeast transformations were washed, plated and incubated for three days at 30 °C in duplicate. After incubation, 36 OD_600nm_ units of cells were harvested and stored at -20 °C prior to plasmid DNA extraction.

### Library sequencing

To determine the change in frequency of variants after selection for GCK activity, we sequenced the GCK ORF before and after growth on glucose medium. The GCK ORF was sequenced in 14 tiles, such that each tile could be sequenced on both strands to reduce base-calling errors. Region one (tile 1-5) and two (tile 6-10) both spanned five tiles, while region three was sequenced in four tiles (tile 10-14).

First, plasmid DNA was extracted from yeast cells for both duplicates of the regional libraries and the GCK WT control pre- and post-selection. Plasmid DNA was extracted from 9 OD_600nm_ units using the ChargeSwitch Plasmid Yeast Mini kit (Invitrogen). Plasmid DNA was adjusted to equal concentrations, and was then used for two rounds of PCR to first amplify each tile and then add index sequences to allow for multiplexing. In the first PCR, each tile was amplified with primers containing a binding site for Illumina sequencing adapters. For each tiling PCR, the following was mixed: 20 µL Phusion High-Fidelity PCR Master Mix with HF Buffer (NEB), 1 µL 10 µM forward primer, 1 µL 10 µM reverse primer, 18 µL plasmid library template. The sequences of forward and reverse primers for each tile are listed in the supplemental material (SKG_tilenumber_fw/rev). Tiles were amplified using the following program: 98 °C 30 sec, 21 cycles of 98 °C 10 sec, 63 °C 30 sec, 72 °C 60 sec, followed by 72 °C 7 min and 4 °C hold.

In the next PCR, Illumina index adapters were added to all amplified tiles. Each PCR consisted of: 20 µL Phusion High-Fidelity PCR Master Mix with HF Buffer (NEB), 2 µL 10 µM i5 indexing adapter, 2 µL 10 µM i7 indexing adapter, 1 µL 1:10 diluted PCR product, 15 µL nuclease-free water. Tiles were amplified using the following program: 98 °C 30 sec, 7 cycles of 98 °C 15 sec, 65 °C 30 sec, 72 °C 120 sec, followed by 72 °C 7 min and hold at 4 °C. An equal volume of each PCR was then pooled and 100 µL were used for gel extraction from a 4% E-Gel EX Agarose Gel (Invitrogen). The fragment size and quality of the extracted DNA were tested using a 2100 Bioanalyzer system (Agilent), and DNA concentration was determined using Qubit (ThermoFisher). Finally, libraries were paired-end sequenced using an Illumina NextSeq 550.

### Sequencing data analysis

The TileSeqMave (https://github.com/jweile/tileseqMave, version 0.6.0.9000) and TileSeq mutation count (https://github.com/RyogaLi/tileseq_mutcount, version 0.5.9) pipelines were used to process sequencing data to obtain variant activity scores.

### Imputation of missing activity scores

The Human Protein Variant Effect Map Imputation Toolkit webserver (Weile et al., 2017; Wu et al., 2019), was used to impute activity scores for missing variants. The webserver was run using standard parameters and with equal quality index on all variant scores. The original and imputed refined scores showed a Pearson’s correlation of 0.985. The imputed scores were only used for Fig. S3 except for the benign variant p.G68D. The imputed score of p.G68D was used for Fig. 3CD and receiver-operating characteristic (ROC) analyses, due to the limited number of benign variants.

### Evolutionary Conservation analysis

The HHblits suite (Remmert et al., 2011; Steinegger et al., 2019) and GEMME package (Laine et al., 2019) were used to evaluate evolutionary distances from the WT GCK sequence (Uniprot ID: P35557 - isoform 1).

The MSA was generated using HHblits with an E-value threshold of 10^-20^ and using UniClust30 as sequence database. The resulting MSA contained 1179 sequences. Two additional filters were applied to the HHblits output MSA: First, all the columns that were not present in the WT GCK sequence were removed. Second, all the sequences with more than 50% gaps were removed, leaving 1079 sequences in the MSA. The GEMME package was run using standard parameters. Median scores were evaluated for each position using all the 19 substitutions.

### Thermodynamic stability measurements (ΔΔG)

Rosetta package (GitHub SHA1 c7009b3115c22daa9efe2805d9d1ebba08426a54) with Cartesian ΔΔG protocol (Frenz et al., 2020; Park et al., 2016) was used to predict changes in thermodynamic stability from the crystal structures (Kamata et al., 2004) of super-open (PDB 1V4T) and closed (PDB 1V4S) conformations of GCK. The values obtained from Rosetta in internal Energy Unit were divided by 2.9 to convert the unit to kcal/mol (Park et al., 2016). Median scores were evaluated for each position using all the 19 substitutions.

### Fasting plasma glucose study population

Variants in GCK were identified using sequencing. Samples were collected from a population-based cohort of 6,058 individuals both with and without diabetes (Glümer et al., 2003), 2,930 patients with newly-diagnosed diabetes (Christensen et al., 2018), patients diagnosed with GCK-MODY (Johansen et al., 2005) and from a population of 1,146 Danish children (Kloppenborg et al., 2018). Individuals were included if they carried one missense GCK variant according to transcript NM_000162 and if a measure of fasting plasma glucose was available. Measures of fasting plasma glucose were examined using a glucose oxidase method (Granutest; Merck, Darmstadt, Germany) in the population based cohort and in samples from patients with known GCK-MODY (Glümer et al., 2003; Johansen et al., 2005), an enzymatic hexokinase method (Gluco-quant Glucose/HK, Roche Diagnostics) in newly diagnosed diabetes patients (Christensen et al., 2018), and using a Dimension Vista® 1500 Analyzer (Siemens, Erlangen, Germany) in children (Kloppenborg et al., 2018). Samples were excluded if fasting plasma glucose level exceeded 9 mM.

### Statistical analyses

Confidence intervals (CIs) were obtained using the SciPy bootstrap function with 10,000 resamples. Statistics associated with activity scores were obtained as documented in the TileSeqMave pipeline (https://github.com/jweile/tileseqMave).

## Acknowledgements

The authors thank Sofie V. Nielsen for helpful conversation on experimental procedures, Jochen Weile who co-developed the TileSeq pipeline used for data analysis, and Amal Al-Chaer for assistance with the Bioanalyzer system and Illumina sequencing. We acknowledge the use of computing resources at the core facility for biocomputing at the Department of Biology, University of Copenhagen.

## Data availability

The Illumina FASTQ files can be accessed at the NCBI Gene Expression Omnibus (GEO) repository under accession number GSE198878. The activity scores have been deposited on MaveDB under accession number tmp:fPGkVi4z7o3jh2l7. All data presented in the manuscript are available in the Supplementary Data file.

## Author contributions

S.G., M.C., M.G., G.S. and A.P.G. performed the experiments. S.G., M.C., M.G., A.G.C., A.P.G., G.S., R.L., D.T., A.S., A.L.G., T.H., F.P.R, K.L.-L. and R.H.-P. analyzed the data. F.P.R., K.L.-L. and R.H.-P. conceived the study. S.G. wrote the paper.

