## Supplementary information for "A comprehensive map of human glucokinase variant activity"

***Supplemental material***

*Fig. S1 (Expression of GCK variants)* *p2*

*Fig. S2 (Co-expression of GKRP and GCK)* *p3*

*Fig. S3 (Imputed map of glucokinase variant activity)* *p4*

*Fig. S4 (Activity scores mapped onto GCK ribbon diagram)* *p5*

*Fig. S5 (Evolutionary analysis of GCK homologous sequences)* *p6*

*Fig. S6 (Correlations between evolutionary conservation and activity scores)* *p7*

*Fig. S7 (Rosetta ΔΔG heatmaps)* *p8*

*Fig. S8 (Positions predicted to shift GCK towards the closed conformation are enriched at the allosteric activator site)* *p9*

*List of primers* *p10*

*References* *p11*

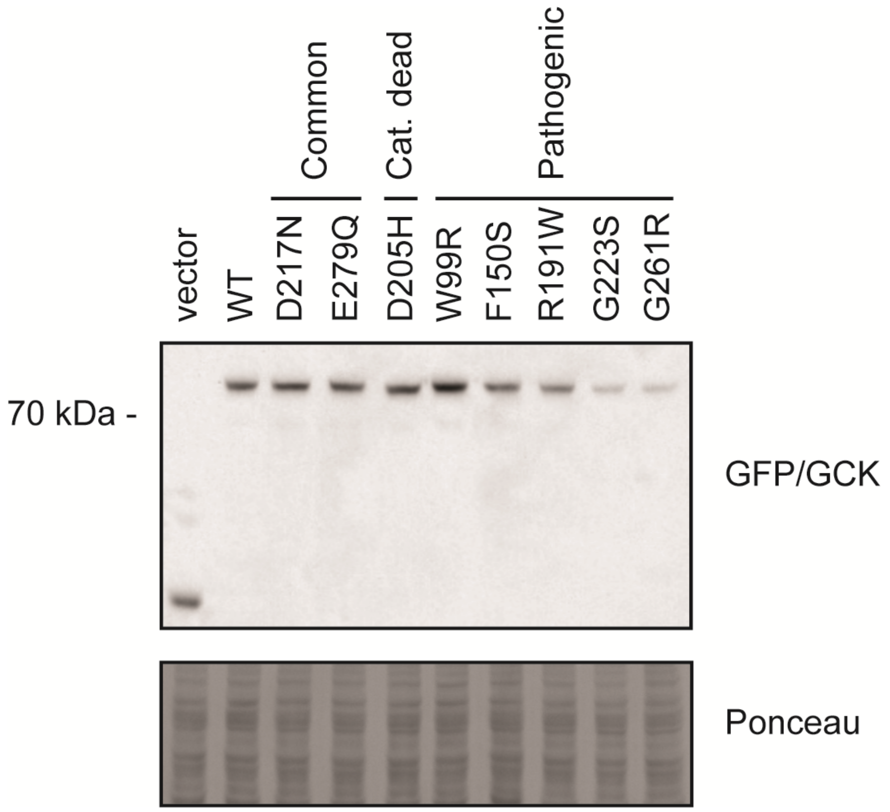


**Figure S1** *Expression of GCK variants*. Western blot showing expression of GCK variants in the *hxk1Δhxk2Δglk1Δ* yeast strain grown in galactose medium.

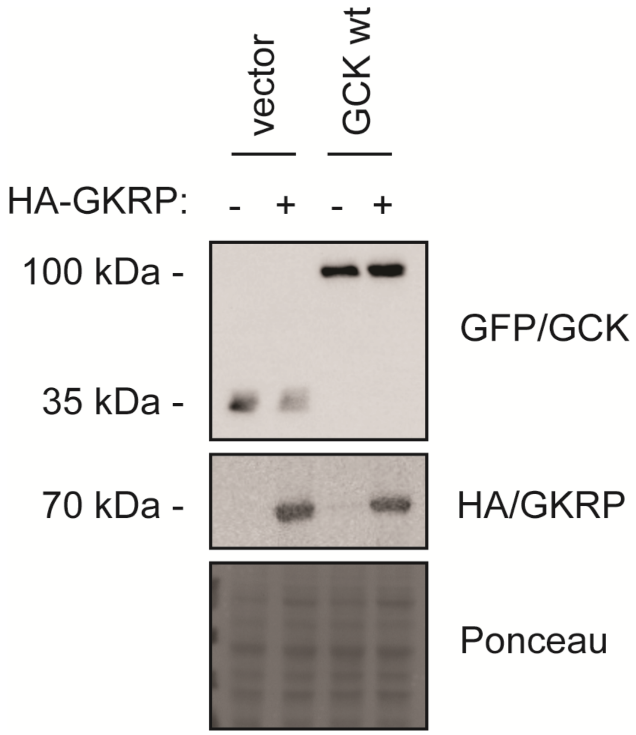


**Figure S2** *Co-expression of GKRP and GCK.* HA-GKRP was co-expressed with either a vector control or wild-type GCK in the *hxk1Δhxk2Δglk1Δ* yeast strain in galactose medium. Protein levels were examined using Western blotting.

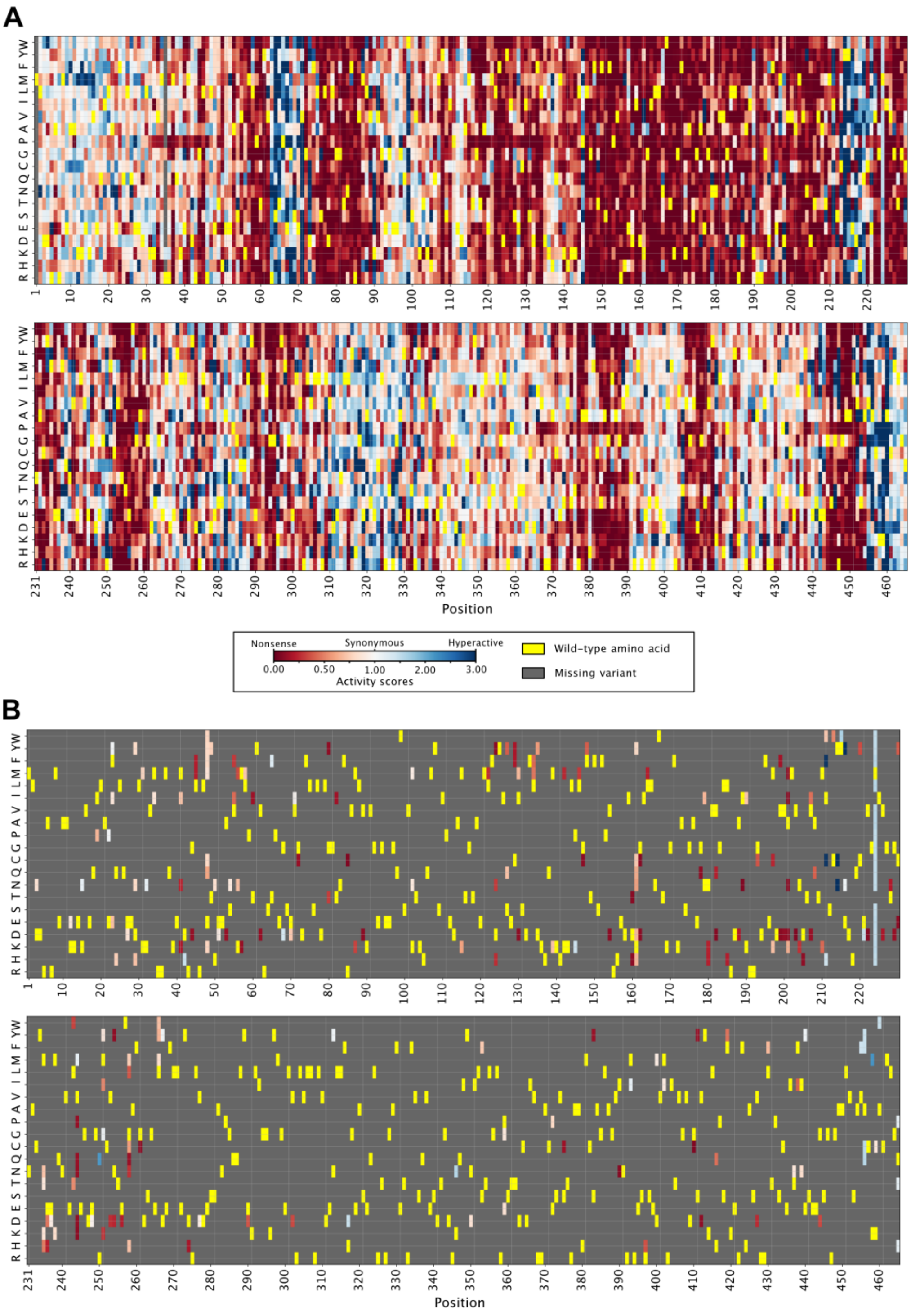


**Figure S3** *Imputed map of glucokinase variant activity*. (A) Map of glucokinase variant activity scores including both experimentally determined and imputed scores. (B) Map of glucokinase variant activity scores showing only imputed scores.

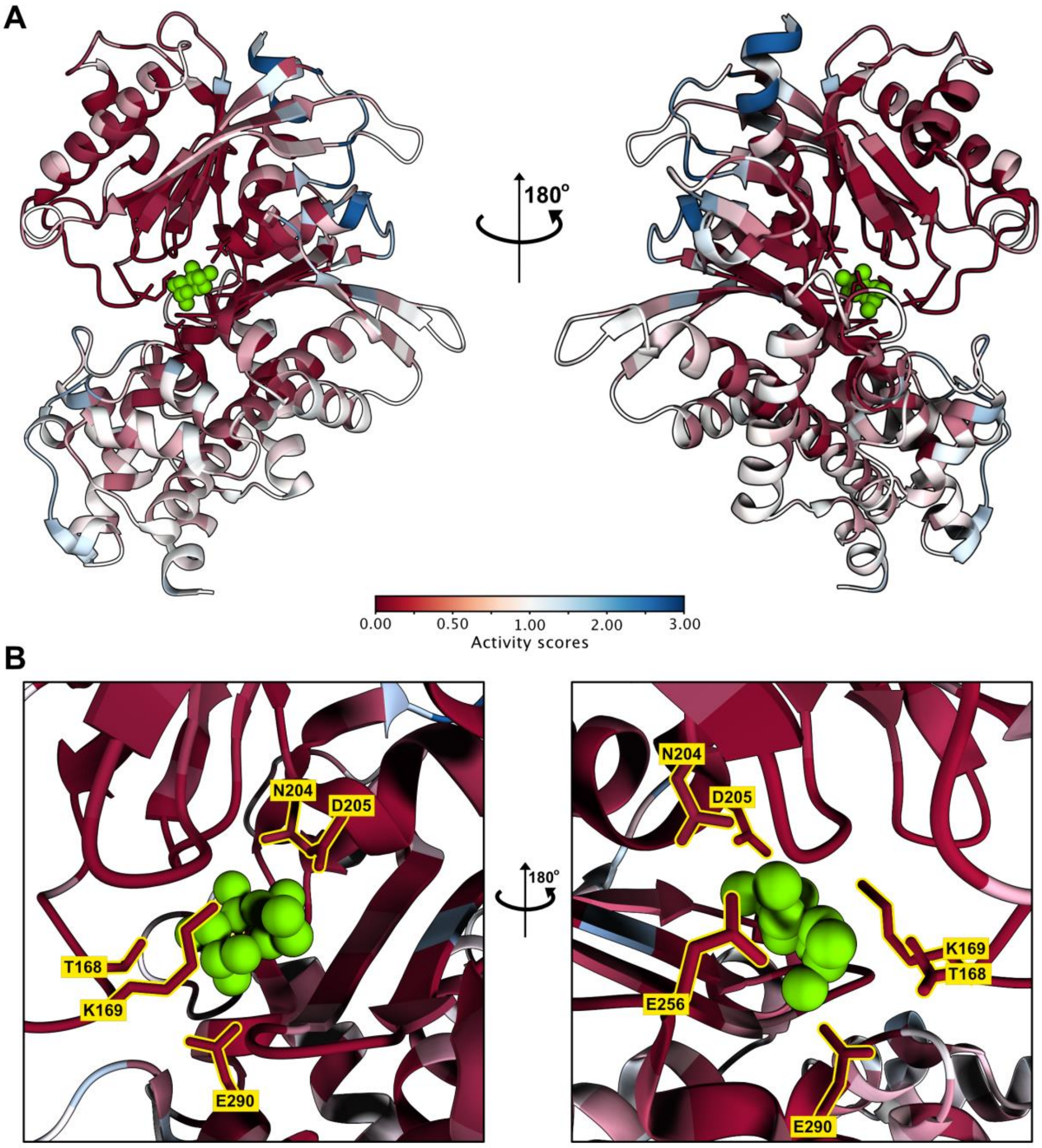


**Figure S4** *Activity scores mapped onto GCK ribbon diagram*. (A) Median activity scores mapped onto glucose-bound GCK (PDB 1V4S). Glucose shown in green. (B) Panels showing the active site from panel A with glucose-binding residues highlighted in yellow and glucose in green.

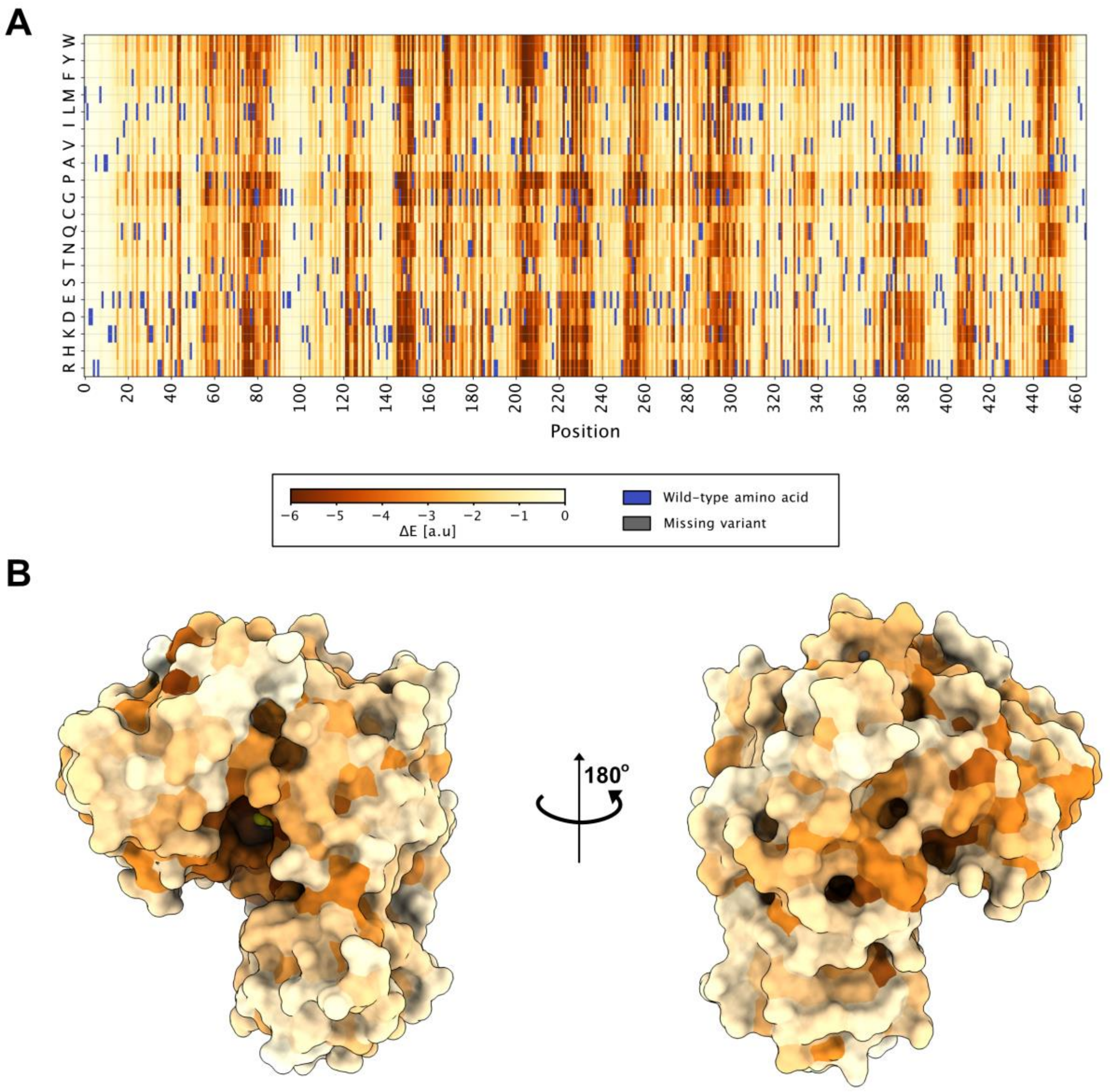


**Figure S5** *Evolutionary analysis of GCK homologous sequences*. (A) Heatmap showing evolutionary conservation analysis of human GCK using a multiple sequence alignment of GCK homologs. The wild-type amino acid at each position is shown in purple. A score close to zero means that a given variant does not have any detrimental effect on GCK function or structural stability, while a high negative score means that a variant affects function or stability. (B) The median evolutionary score for each position is mapped onto the closed conformation of GCK (PDB 1V4S), using the same color scheme as in A.

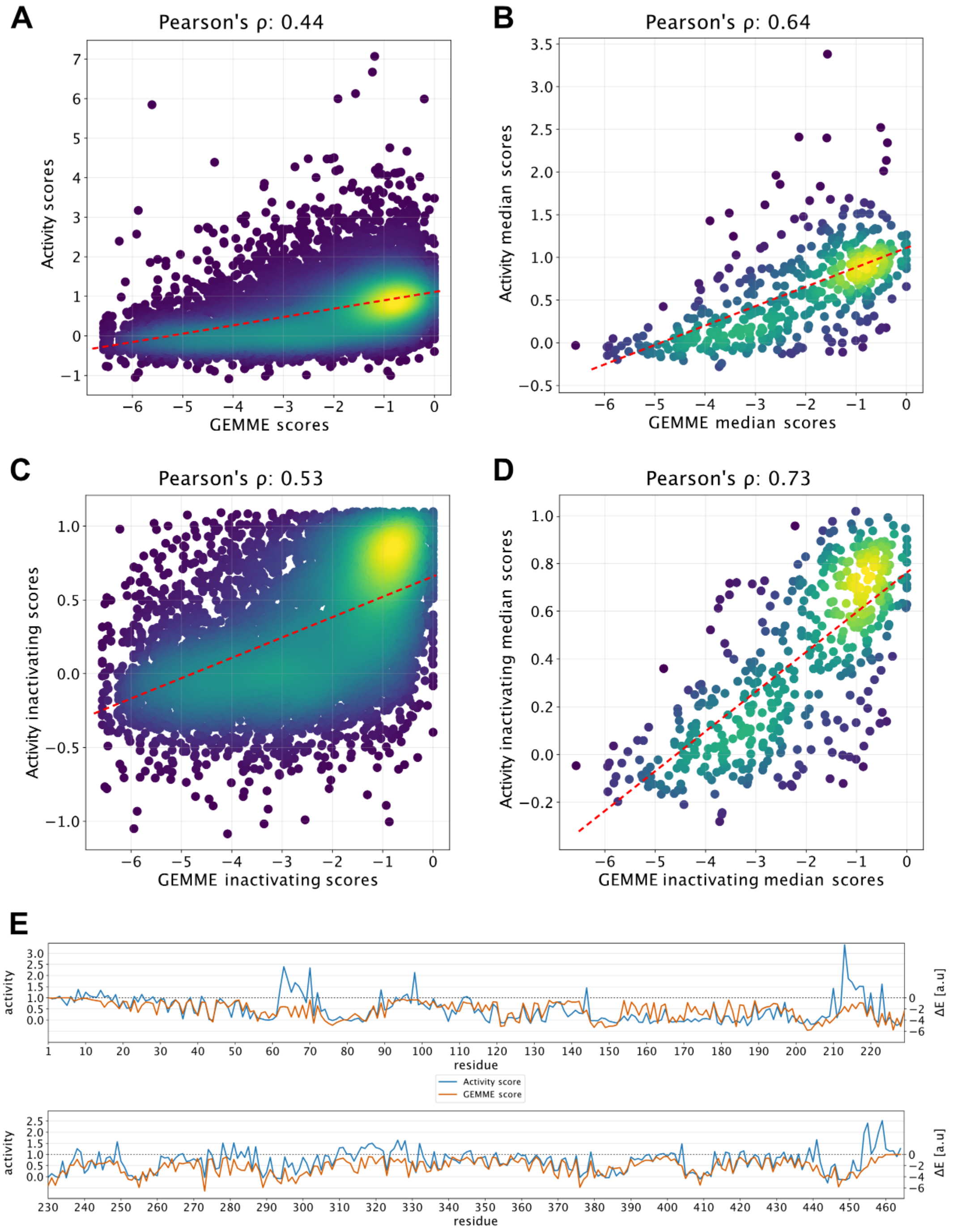


**Figure S6** *Correlations between evolutionary conservation and activity scores.* (A) Correlation between GEMME scores and activity scores for all variants. The dotted red line shows the best fitting curve. (B) Correlation between GEMME scores and activity scores using the median score at each position. The dotted red line indicates the best fitting curve. (C) The correlation from panel A with hyperactive variants (activity score > 1.18) excluded. (D) The correlation from panel B with hyperactive variants (activity score > 1.18) excluded. (E) Line plot of the median activity and GEMME score at each position along the GCK sequence.

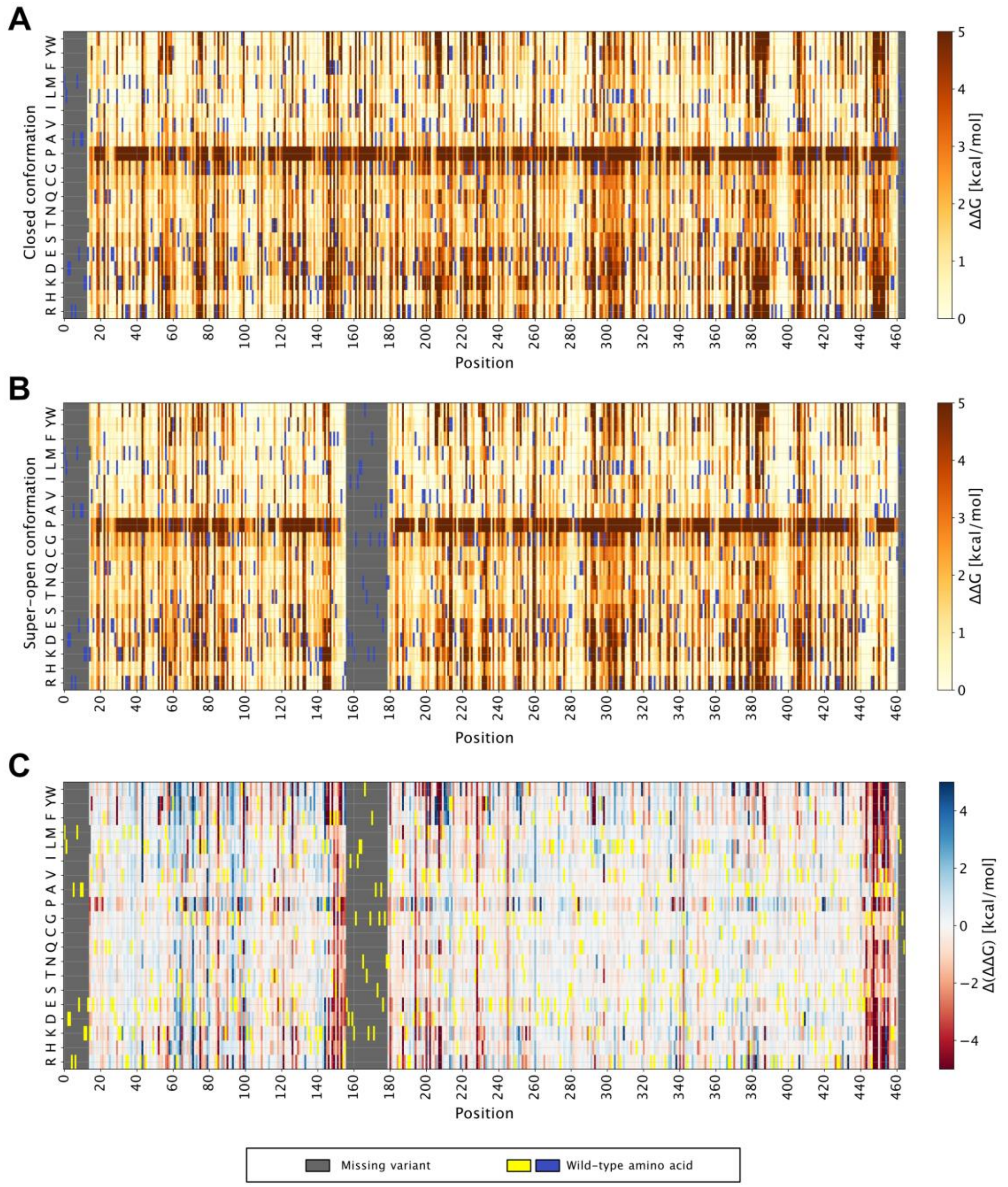


**Figure S7** *Rosetta ΔΔG heatmaps.* (A) Heatmap showing ΔΔG scores for each single variant in the closed conformation (PDB 1V4S). A score close to zero means that a given variant has a wild-type-like stability, while a positive score means that a variant is less stable than WT. (B) Heatmap showing ΔΔG scores for each single variant in the super-open conformation (PDB 1V4T) with variant scores the same as in panel A. (C) Heatmap showing the difference between ΔΔG in the closed and super-open conformation for each variant. Variants with a positive score (blue shades) destabilize the super-open conformation relative to the closed state. Conversely, variants with a negative score (red shades) destabilize the closed conformation relative to the super-open state.

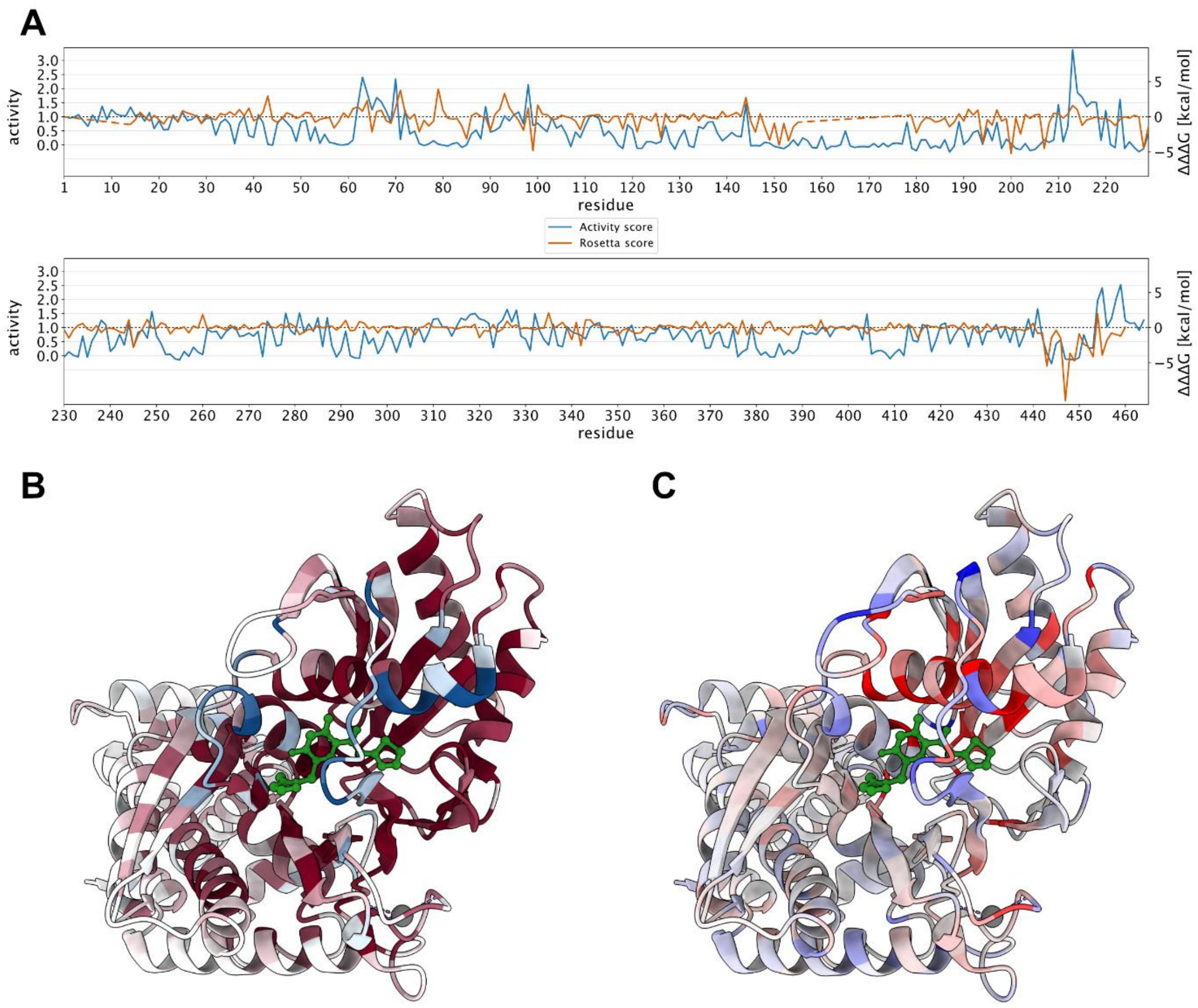


**Figure S8** *Positions predicted to shift GCK towards the closed conformation are enriched at the allosteric activator site.* (A) Line plot of median activity and Rosetta score (ΔΔΔG) at each position along the GCK sequence. (B) Structure of GCK bound to the synthetic allosteric activator compound A (PDB 1V4S) colored by median activity score. Compound A binds to the allosteric activator site (Kamata et al., 2004) and is shown in green. (C) Structure of GCK bound to compound A (PDB 1V4S) colored by ΔΔΔG. Compound A shown in green.

**List of primers**

SKG_1: GCATGCCAATACTGGTAGATCACCGTAAAACGACGGCCAGTCTTAA

SKG_2: GCATGCCAATACTGGTAGATCACC

SKG_3: CAGGAAACAGCTATGACCATGT

SKG_4: GGGGACAAGTTTGTACAAAAAAGCAGGCTTCGC

SKG_5: GGGGACCACTTTGTACAAGAAAGCTGGGT

SKG_tile1_fw: TACACGACGCTCTTCCGATCTGCAGGCTTCGCCACC

SKG_tile1_rev: AGACGTGTGCTCTTCCGATCTTCAAACCTCTATCCATTTCCTTTTG

SKG_tile2_fw: TACACGACGCTCTTCCGATCTGATTTGAAGAAAGTTATGAGAAGAATG

SKG_tile2_rev: AGACGTGTGCTCTTCCGATCTAATCTAATGACAAAAAGTCACCAACTT

SKG_tile3_fw: TACACGACGCTCTTCCGATCTTTAGATCTACACCAGAAGGTTCAG

SKG_tile3_rev: AGACGTGTGCTCTTCCGATCTTCTGGGATTGAGTACATTTGATGCT

SKG_tile4_fw: TACACGACGCTCTTCCGATCTAAGGTCAATGGTCTGTTAAGACAA

SKG_tile4_rev: AGACGTGTGCTCTTCCGATCTAATGGCAATTTCTTATGCTTCATTTGA

SKG_tile5_fw: TACACGACGCTCTTCCGATCTATGTATCTCAGATTTCTTGGATAAGCA

SKG_tile5_rev: AGACGTGTGCTCTTCCGATCTACCTTCTGCACCTGAAGCT

SKG_tile6_fw: TACACGACGCTCTTCCGATCTGTTGAACTGGACAAAGGGTTTTAA

SKG_tile6_rev: AGACGTGTGCTCTTCCGATCTGTAACAAGAGATCATTGTTGCAACA

SKG_tile7_fw: TACACGACGCTCTTCCGATCTGGATGTTGTTGCTATGGTTAACGATAC

SKG_tile7_rev: AGACGTGTGCTCTTCCGATCTGTCACCTTCAACCAATTCAACG

SKG_tile8_fw: TACACGACGCTCTTCCGATCTCGCATGTTACATGGAAGAAATGCAAAA

SKG_tile8_rev: AGACGTGTGCTCTTCCGATCTTGAAGATTCATCAACCAATCTATCGTA

SKG_tile9_fw: TACACGACGCTCTTCCGATCTGGTGAATTGGATGAATTCTTGTTGGAA

SKG_tile9_rev: AGACGTGTGCTCTTCCGATCTAATTTTCATCAACCAATCTCAACAAAA

SKG_tile10_fw: TACACGACGCTCTTCCGATCTAATACATGGGTGAATTAGTTAGATTGG

SKG_tile10_rev: AGACGTGTGCTCTTCCGATCTTTTCTGTCACCAGTATCTGATTCAA

SKG_tile11_fw: TACACGACGCTCTTCCGATCTCATTCGAAACAAGATTCGTTTCTCAAG

SKG_tile11_rev: AGACGTGTGCTCTTCCGATCTGCAGCTCTAGTTGAAACAGATTCAC

SKG_tile12_fw: TACACGACGCTCTTCCGATCTCAGATTGTGATATTGTTAGAAGAGCTT

SKG_tile12_rev: AGACGTGTGCTCTTCCGATCTACAGAACCATCAACACCAACTG

SKG_tile13_fw: TACACGACGCTCTTCCGATCTCTAGATCAGAAGATGTTATGAGAATCA

SKG_tile13_rev: AGACGTGTGCTCTTCCGATCTCCTGAACCTTCTTCAGATTCAATAAAA

SKG_tile14_fw: TACACGACGCTCTTCCGATCTAAGATTGACTCCATCATGTGAAATCAC

SKG_tile14_rev: AGACGTGTGCTCTTCCGATCTTCTTATAATGCCAACTTTGTACAAGAA
